# Genetic background modifies CNS-mediated sensorimotor decline in the AD-BXD mouse population

**DOI:** 10.1101/657122

**Authors:** Kristen M.S. O’Connell, Andrew R. Ouellette, Sarah M. Neuner, Amy R. Dunn, Catherine C. Kaczorowski

**Author notes:** Co-senior authors (Kristen M.S. O’Connell, Catherine C. Kaczorowski).

## Abstract

Many patients with Alzheimer’s dementia also exhibit non-cognitive symptoms such as sensorimotor deficits, which can precede the onset of the hallmark cognitive deficits and significantly impact daily activities and an individual’s ability to live independently. However, the mechanisms underlying sensorimotor dysfunction in AD and the relationship between sensorimotor and cognitive decline remain poorly understood, due in part to a lack of translationally relevant animal models. To address this, we recently developed a novel model of genetic diversity in Alzheimer’s disease, the AD-BXD genetic reference panel. In this study, we investigated sensorimotor deficits in the AD-BXDs and their relationship to cognitive decline in these mice. We found that both age- and AD-related declines in coordination, balance, and vestibular function vary significantly across the panel, indicating that genetic background strongly influences the expressivity of the familial AD mutations used in the AD-BXD panel and their impact on motor function. Further, we found that motor decline is not associated withcognitive decline in either AD or aging, suggesting that sensorimotor deficits in AD occur – in part - through distinct mechanisms. Overall, the results presented here suggest that AD-related sensorimotor decline is strongly dependent on background genetics and is independent of dementia and cognitive deficits, suggesting that effective therapeutics for the entire spectrum of AD symptoms will likely require interventions targeting each distinct domain involved in the disease.

## Introduction

Alzheimer’s dementia (AD) is defined by the slow progression of cognitive deficits, including memory loss and dementia, accompanied by the accumulation of β-amyloid (Aβ) plaques and hyperphosphorylated tau tangles (Selkoe, 1991). However, patients with AD often experience additional non-cognitive symptoms that significantly impact daily life for both patients and caregivers that lead to an inability to live independently and a need for long-term care (Albers et al., 2015; Buchman and Bennett, 2011; Portet et al., 2009; Scarmeas et al., 2004; Voglein et al., 2019). Among these non-cognitive symptoms are deficits in sensorimotor function such as gait slowing, loss of balance and coordination, sarcopenia and muscle weakness, and increased frailty. Furthermore, the emergence of these symptoms during the preclinical stage of AD is associated with worse AD-related cognitive decline than in individuals who do not exhibit motor-related symptoms. (Portet et al., 2009).

Development of motor deficits prior to the hallmark memory loss associated with AD (Albers et al., 2015; Buchman and Bennett, 2011) suggests that they may represent some of the very earliest events in the pathogenesis of AD. Unfortunately, motor dysfunction and other non-cognitive symptoms in AD are relatively poorly understood compared to the cognitive deficits and memory loss so closely associated with the disease. It therefore remains unknown whether motor impairment represents an early biomarker of disease, or if it is part of a chain of causal events leading to dementia. It is essential to determine whether the association between AD-related cognitive decline and motor function shares a common mechanism, if shared pathogenesis exists, understanding the causal factors underlying motor symptoms may enhance our ability to identify new pathways for therapeutics that address both cognitive and motor domains. Alternatively, if they have distinct causal mechanisms, then therapies targeting cognitive symptoms may be ineffective in treating motor dysfunction and thus, fail to prevent or delay the loss or independence or the need for long-term institutional care.

Significant progress in understanding the pathophysiology of AD has been made by utilizing mouse models incorporating familial mutations in amyloid precursor protein (APP) and/or presenilin-1 (PSEN1), originally designed to recapitulate the cognitive symptoms of AD. Notably, several of these models have been reported to also exhibit sensorimotor deficits, suggesting that motor impairments are an inherent part of the disease process (Jawhar et al., 2012; O’Leary et al., 2018a; O’Leary et al., 2018b; O’Leary et al., 2018c; Seo et al., 2010; Wirths and Bayer, 2008; Wirths et al., 2008). However, infrequent assessment of motor phenotypes in AD animal models, variability in the tests used to assess motor function, and use of a single sex has led to conflicting reports on the impact of AD transgenes on motor function (Jawhar et al., 2012; Lalonde et al., 2004; O’Leary et al., 2018c; Seo et al., 2010; Stover et al., 2015).

The etiology of AD in humans is complex and although age is the greatest risk factor for developing AD, it is increasingly clear that genetics and family history play a significant role (Gatz et al., 1997; Ryman et al., 2014). Most animal models are developed on single or a few inbred backgrounds (Onos et al., 2016), with little or no genetic variation, presenting a challenge for identifying and investigating AD symptoms such as motor dysfunction, which can vary considerably in its presentation in human populations. As such, a major barrier to understanding the overlap (or lack thereof) of mechanisms underlying motor dysfunction and cognitive symptoms in AD is a lack of translationally relevant animal models that reflect individual differences in symptom onset and progression in the human population.

To address this, we recently developed the AD-BXD panel (Neuner et al., 2019a), which combines the well-characterized 5XFAD model of AD (Oakley et al., 2006) with the BXD genetic reference panel (Peirce et al., 2004; Taylor et al., 1999) to create a novel AD model that incorporates both causal AD mutations and naturally-occuring genetic diversity to better model the human disease. In the present study, we assess the impact of AD, normal aging, and naturally occurring genetic variation on sensorimotor-related phenotypes and their relationship to cognitive outcomes using the AD-BXD panel. Because the panel also includes the non-transgenic controls, we can distinguish the influence of familial AD mutations from the normal decline in motor skills commonly observed in aging. We hypothesized that the AD-BXD panel would exhibit age-related decline in sensorimotor function that is exacerbated by the presence of the AD transgene and that diverse genetic backgrounds would influence the expressivity of the 5XFAD transgene to modify the onset and severity of motor-related phenotypes.

## Results

### Balance and motor coordination are impaired in AD-BXD mice in a genetic-background dependent manner

Since balance and coordination skills are often impaired in human AD patients (Buchman and Bennett, 2011), we assessed these sensorimotor domains in the AD-BXD panel using the narrow beam task, which is a well-characterized and validated assay for balance and coordination in mice (Brooks and Dunnett, 2009; Carter et al., 2001). To determine the impact of genetic background on AD- and age-related impairments in these domains, we measured performance on the narrow beam apparatus in 27 strains of AD-BXD mice (27 strains female, 18 strains male) at 6 and 14 months (m) of age. We also tested age-matched Ntg-BXD strains to assess the impact of normal aging on this task.

As shown in Figure 1A, as a population, AD-BXD mice took significantly longer to cross the beam compared to the non-carrier controls (Genotype: F_1,886_ = 24.5, p = 9.08 × 10^−7^). There was a main effect of age on narrow beam performance (6m v 14m: F_1, 886_ = 3.9, p = 0.047) and presence of the AD transgene significantly exacerbated age-related decline (Age*Genotype: F_1,886_ = 7.6, p = 0.005) (Figure 1B). Post hoc comparisons using the Tukey HSD test indicated that this effect was primarily driven by the presence of the AD transgene and was not due to decline due to normal aging, as there was no significant effect of age in the Ntg-BXD (6 m-14 m, adj. p = 0.67). Furthermore, although there was a trend for 6m AD-BXD mice to perform worse on this task than their age-matched Ntg-BXD controls, this difference was not significant (adj. p = 0.081). Lastly, although there was no significant main effect of sex on narrow beam performance across the panel (Sex: F_(1, 886)_ = 0.052, p = 0.81), there were significant interactions between both age and sex (Age*Sex: F_(1, 886)_ = 9.16, p = 0.002) and sex and genotype (Sex*Genotype: F_(1, 886)_ = 7.9, p = 0.0045), with post hoc comparisons suggesting that impairment due to age and AD status was greater in females than males (Figure 1C).

**Figure 1:**
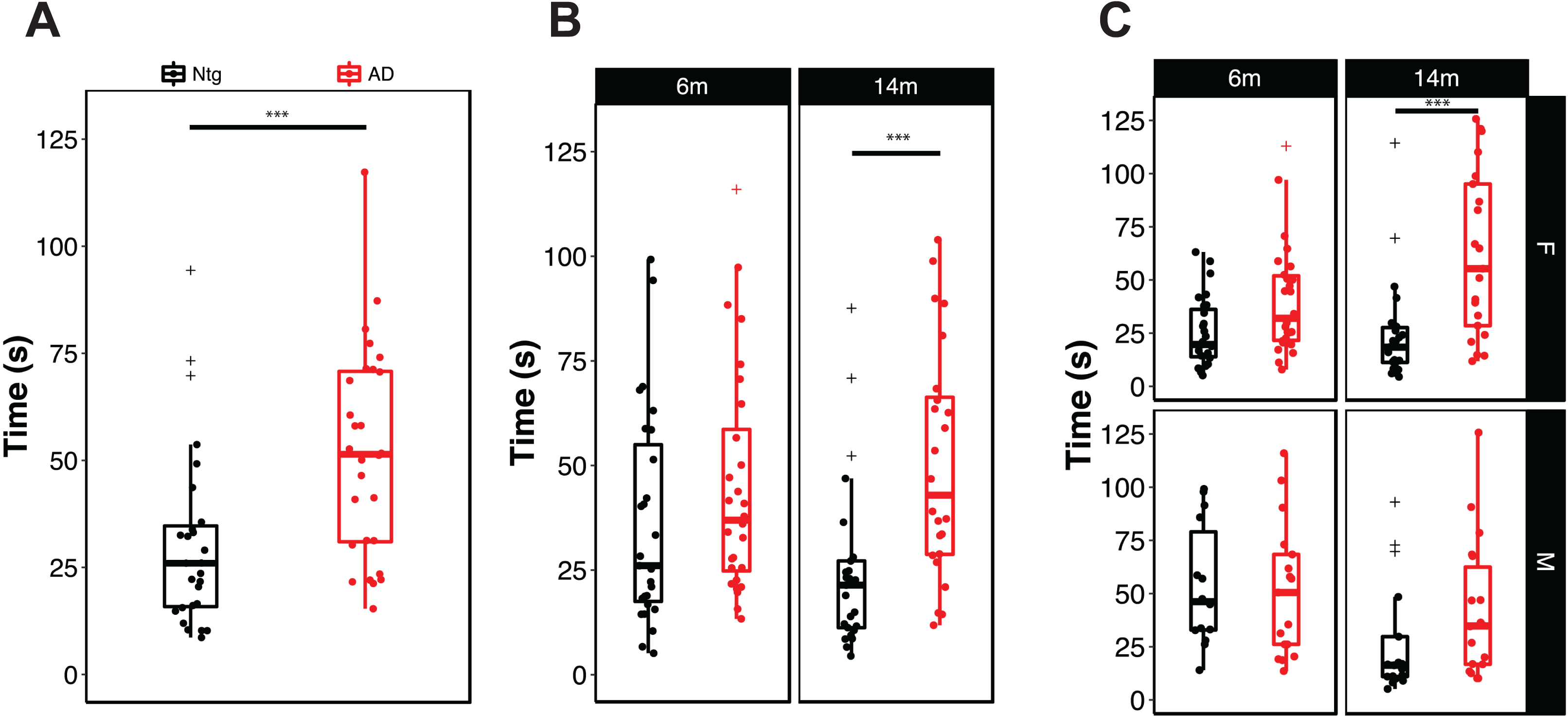
Presence of the high-risk AD transgene impairs motor coordination and balance in an age- and sex-dependent manner. **A)** Mice carrying the 5XFAD transgene take significantly longer to cross the narrow beam apparatus, indicating impaired balance and coordination. **B)** Stratification of narrow beam performance by age and **C)** by age and sex. Each point represents a strain average and statistical outliers are indicated by a + sign. *** = adj. p <0.0001 following post hoc testing using Tukey’s HSD.

To assess the influence of genetic background on balance and coordination, we compared narrow beam performance across the Ntg- and AD-BXD strains phenotyped in this study. Figure 2 illustrates the average time taken to cross the narrow beam by strain at 6m (Figure 2A) and 14m (Figure 2B) in the Ntg-BXDs (*left*) and AD-BXDs (*right*). There is a main effect of genetic background on the time taken to cross the narrow beam in the the Ntg- and AD-BXD panels (Strain_Ntg_: F_(26, 363)_ = 3.35, p = 1.6 × 10^−7^ Strain _AD_: F_27,474_) = 2.7, p = 1.02 × 10^−5^), suggesting that naturally occurring genetic variation itself strongly contributes to performance on the narrow beam task in both genotypes. The impact of age on narrow beam performance is illustrated in Figure 2C by plotting the difference in time to cross at 14m from 6m in the Ntg (*left)* and AD (*right*) BXD strains, demonstrating that overall, very few Ntg-BXD strains exhibit age-related decline the narrow beam performance, while age-related decline is exacerbated in most (but not all) AD-BXD strains. Notably, although there are significant main effects of both strain and genotype as described above, there is no significant correlation between the Ntg- and AD-BXD strains on narrow beam performance in either males (*r* = 0.24, p = 0.35) or females (*r* = 0.27, p = 0.21) (Supplemental Figure 1), suggesting that genotype*strain interactions have a greater impact on narrow beam performance than strain alone. Consistent with the pronounced influence of background strain, heritability 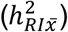 estimates comparing the between-strain variance (an estimate of variation due to genetic factors) to total sample variance (an estimate of variation due to environmental, technical, and genetic factors) indicate that genetics significantly influence phenotypic variation (Table 1).

**Table 1:** Heritability estimates for sensorimotor traits in Ntg- and AD-BXD strains. Heritability 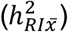 was calculated as the ratio of between-strain variance (i.e., trait variance due to genetic factors) to total variance (i.e., variance due to technical and environmental factors, assessed as within-strain variance, plus genetic variance) normalized to the average number of biological replicates per strain (*n*) according to (B<u>elknap, 1998).</u>

| **Ntg-BXD** |  |  |  |  |
| --- | --- | --- | --- | --- |
| **Trait** | **Between-strain Variability** | **Avg. Within-strain variability** | **Avg. *n*/strain** | **$h^2_{RI\bar{x}}$** |
| **6m Narrow Beam** | 667.3893 | 2351.59 | 8.68 | 0.7 |
| **14m Narrow Beam** | 414.7125 | 1428.3 | 7.39 | 0.7 |
| **6m Inclined Screen** | 63.05047 | 147.417 | 8.68 | 0.8 |
| **14m Inclined Screen** | 92.87501 | 425.05 | 7.39 | 0.6 |
| **6m Grip Strength** | 0.08793 | 0.188 | 8.68 | 0.8 |
| **14m Grip Strength** | 0.074671 | 0.179 | 7.39 | 0.8 |
| **AD-BXD** |  |  |  |  |
| **Trait** | **Between-strain Variability** | **Avg. Within-strain variability** | **Avg. *n*/strain** | **$h^2_{RI\bar{x}}$** |
| **6m Narrow Beam** | 734.1014 | 2697.44 | 10.77 | 0.7 |
| **14m Narrow Beam** | 1357.38 | 4261.44 | 9.125 | 0.7 |
| **6m Inclined Screen** | 90.23455 | 361.28 | 10.77 | 0.7 |
| **14m Inclined Screen** | 307.7143 | 1362.3 | 9.125 | 0.7 |
| **6m Grip Strength** | 0.135841 | 0.194 | 10.77 | 0.9 |
| **14m Grip Strength** | 0.075719 | 0.211 | 9.125 | 0.8 |

**Figure 2:**
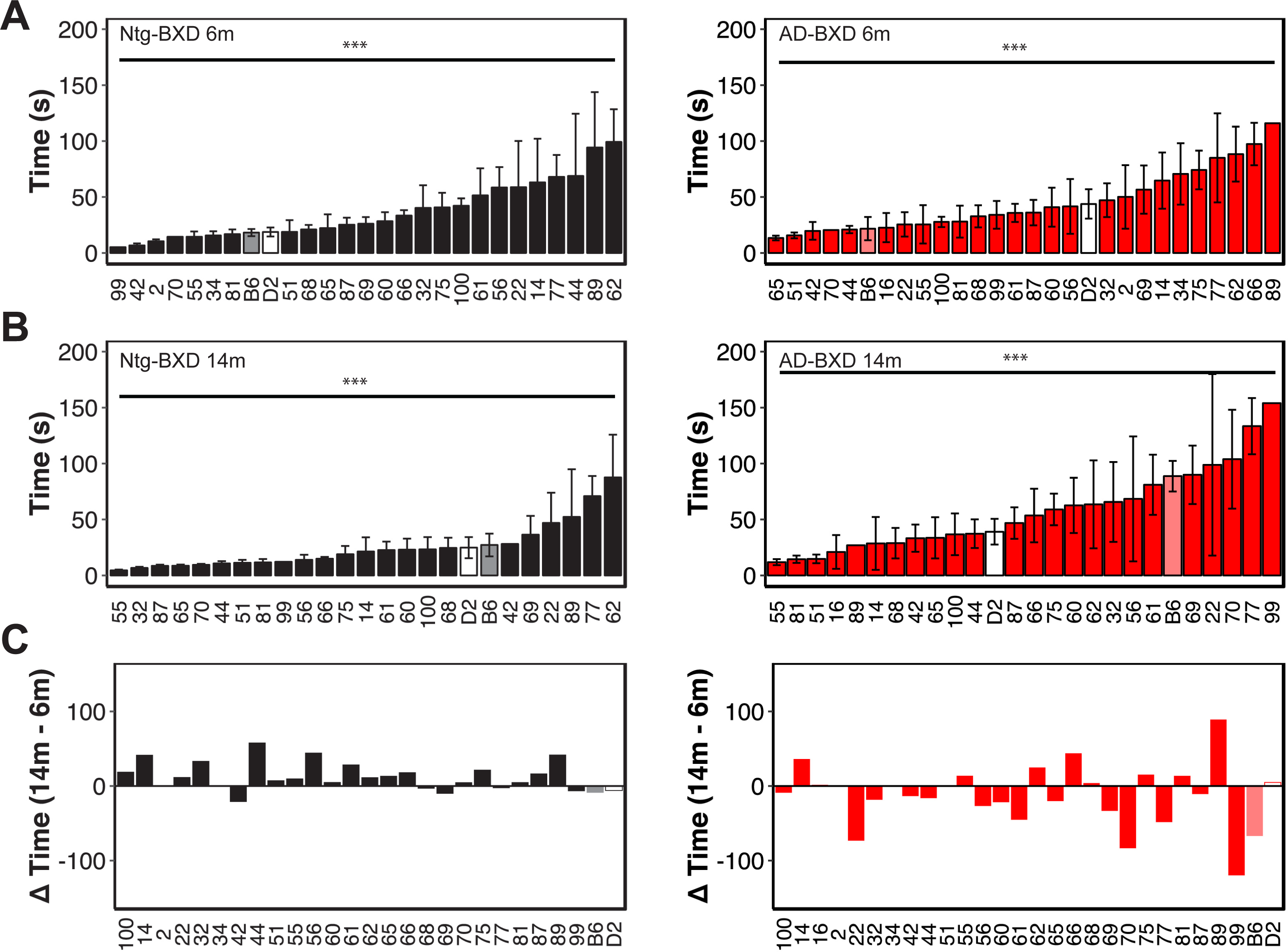
Genetic background influences motor coordination and balance on the narrow beam task in both normal aging and AD. **A)** Mean time to cross the narrow beam averaged by strain in 6m old Ntg-BXD (*left*) and AD-BXD (*right*) strains. **B)** Mean time to cross the narrow beam averaged by strain in 14m old Ntg-BXD (*left*) and AD-BXD (*right*) strains. **C)** Average age-related decline by strain in the Ntg-BXD (*left*) and AD-BXD (*right*) panels. Decline was calculated by subtracting performance at 6m of age from that measured at 14m of age. Data are presented as mean ± SEM. In all panels, number on the x-axis indicates the BXD strain used to generate each line. *** = p <0.0001.

The AD-BXD panel exhibits significant strain-dependent cognitive decline (Neuner et al., 2019a), so to determine whether cognitive performance in this panel correlates with sensorimotor ability, we assessed the relationship between narrow beam performance and memory as measured using the contextual fear memory task (Neuner et al., 2019a). We found no significant correlation between motor performance and either short-term memory (contextual fear acquisition, CFA) or long-term memory (contextual fear memory, CFM; from (Neuner et al., 2019a)) in either Ntg- or AD-BXD mice of either sex (Supplemental Figure 1A), suggesting that the mechanism(s) underliying deficits in balance and coordination are, in part, unrelated to aging or AD-related cognitive impairment in this panel of mice.

### AD-related impairments in motor coordination and vestibular function

Human AD patients often exhibit impairments in balance and orientation suggestive of deficits in vestibular function and proprioception, domains that are assessed in mice using the inclined screen test. In this test, mice are placed head down on a wire mesh screen fixed at a 45° angle and their natural reflex to reposition themselves head up measured, with longer righting times suggestive of impairments in proprioceptive and vestibular systems (Fan et al., 2010). There was a significant effect of AD genotype on this task (Figure 3A), with AD-BXD mice requiring significantly longer time to reorient themselves than Ntg-BXD mice (Genotype: F_(1,886)_ = 55.21, p = 2.5× 10^−13^). There was also a significant effect of age on inclined screen performance (Figure 3B), with 14m old mice taking significantly longer than 6m old mice (Age: F_(1,886)_ = 59.8, p = 2.77 × 10^−14^). For this task, there was a significant interaction between age and genotype (Age*Genotype: F_(1,886)_ = 20.33, p = 7.3 × 10^−6^). As with the narrow beam task, post hoc multiple comparisons indicated that Ntg-BXD mice do not exhibit significant age-related decline (adj. p = 0.19), while incline screen performance is significantly worsened with age in the AD-BXD mice (adj. p <0.000001), suggesting a strong interaction between age and genotype on vestibular function and proprioception. Again, there was no main effect of sex on inclined screen performance (Sex: F_(1,886)_ = 1.5, p = 0.2), nor was there a significant interaction between sex and age (Age*Sex: F_(1,886)_ = 1.14, p = 0.28). There was an interaction between sex and genotype (Genotype*Sex: F_(1,886)_ = 3.87, p = 0.049), which post hoc comparisons indicate is due to worse performance by both male and female AD mice relative to their Ntg counterparts (Male adj. p = 0.001; female adj. p < 0.000001, Figure 3C).

**Figure 3:**
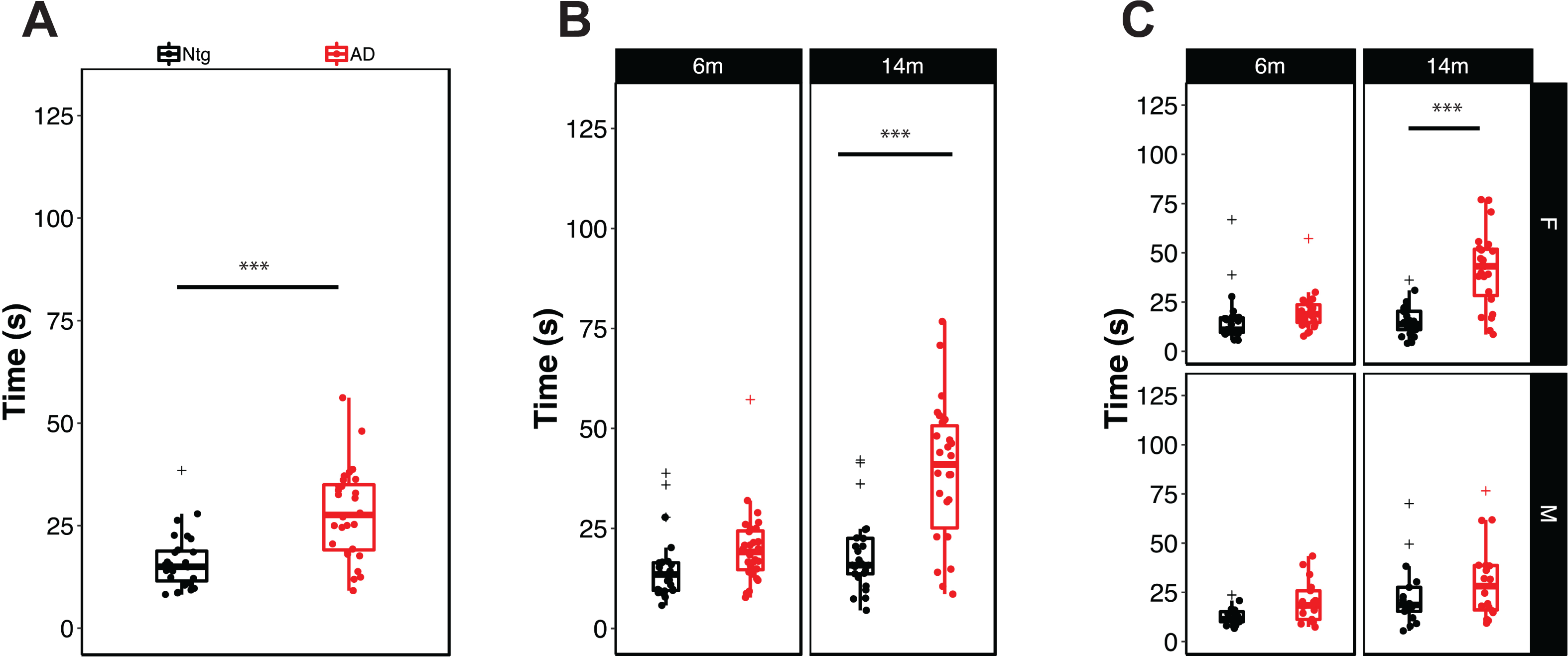
Impairments in motor coordination, vestibular function, and proprioception are exacerbated by the AD transgene. **A)** Mice carrying the 5XFAD transgene take significantly longer to right themselves on the inclined screen apparatus, indicating impaired proprioception and vestibular function. **B)** Stratification of inclined screen performance by age and **C)** by age and sex. Each point represents a strain average and statistical outliers are indicated by a + sign. *** = adj. p<0.0001 following post hoc testing using Tukey’s HSD.

To assess the impact of genetic variation on inclined screen performance, we plotted righting latency as a function of strain average (Figure 4). Analysis of individual strain performance on this task demonstrated a significant main effect of strain in both Ntg- and AD-BXD strains (Strain_Ntg_: F_(26,339)_ = 2.04, p = 0.002; Strain_AD_: F_(27, 474)_ = 2.2, p = 0.0004), once again suggesting a strong influence of naturally occurring genetic variation on vestibular function and proprioception (Figure 4A). On this task, there was a no significant interaction between strain and age in either genotype (Age*Strain_Ntg_: F_(24,339)_ = 0.79, p = 0.74; Age*Strain_AD_: F_(25, 450)_ = 1.2, p = 0.22). We found a significant interaction between strain, sex, and genotype, indicating that female mice from AD strains are impaired relative to male AD strains, which may account for the slight difference in decline between 6m and 14m of age shown in Figure 4C. As with narrow beam, there was no significant correlation between inclined screen performance in Ntg-BXD strains compared with AD-BXD strains (females: *r* = 0.21, p = 0.34; males: *r* = - 0.13, p = 0.62) (Supplemental figure 1B), suggesting that expressivity of the transgene is not equal across strains, but instead depends on genetic background. As with narrow beam, 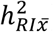 estimates indicate that genetic background accounts for much of the variation in inclined screen performance (Table 1).

**Figure 4:**
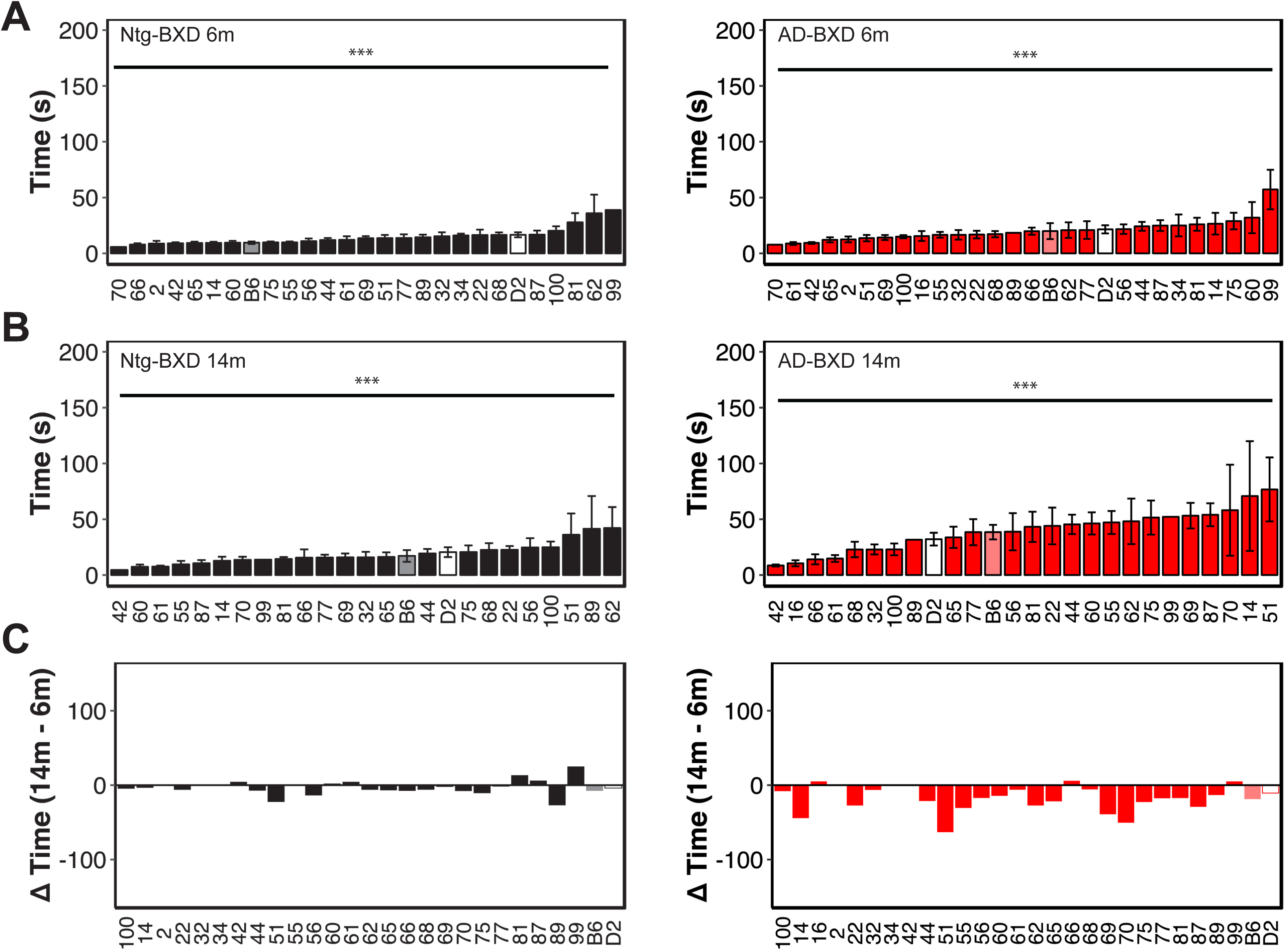
Genetic background influences vestibular function and proprioception on the inclined screen task in both normal aging and AD. **A)** Mean righting time on the inclined screen averaged by strain in 6m old Ntg-BXD (*left*) and AD-BXD (*right*) strains. **B)** Mean righting time on the inclined screen average by strain in 14m old Ntg-BXD and AD-BXD strains. **C)** Average age-related decline by strain in the Ntg-BXD (*left*) and AD-BXD (*right*) panels. Decline was calculated by subtracting performance at 6m of age from that measured at 14m of age. Data are presented as mean ± SEM. In all panels, number on the x-axis indicates the BXD strain used to generate each line. *** = p <0.0001.

Similar to the narrow beam task, we found no correlation between performance on the inclined screen test and long-term CFM in male Ntg- and AD-BXD strains or female AD-BXD strains (Supplemental Figure 3), suggesting impairments in vestibular function and proprioception may be independent of cognitive decline. Interestingly, we did observe a significant positive correlation between inclined screen performance and long-term CFM in female Ntg-BXD strains (Supplemental figure 3B, *left*); however, it should be noted that there is very little phenotypic variation in the inclined screen task compared to CFM, with all female Ntg-BXD strains performing very well on this task.

### Muscle strength decreases with age in both Ntg- and AD-BXD mice

Sarcopenia and muscle weakness are common in aging humans and grip strength is frequently used to monitor strength in the aging population. As in humans, grip strength can be similarly readily assessed in mice and can reveal declines in muscle mass and strength in aging mice. To assess grip strength, mice were allowed to grasp the grip strength apparatus with all four paws while being suspended by the tail. Muscle strength in the fore- and hindlimbs was measured by pulling the mouse away from the bar by the tail to determine the force exerted during the animal’s attempt to maintain its grip on the bar; greater force is indicated by more negative values. As shown in Figure 5A, there was no significant effect of genotype on grip strength, with Ntg and AD mice performing nearly equally (Genotype: F_1,892)_ = 0.04, p = 0.83). As expected, there was a significant effect of age on grip strength, with both Ntg and AD mice exhibiting significant decline with age from 6m to 14m (Age: F_(1,892)_ = 1444.70, p < 2.2 × 10^−16^) (Figure 5B) and female mice exhibiting lower grip strength than male mice (Sex: F_(1,892)_ = 40.97, p = 3.03 × 10^−10^) (Figure 5C). However, there was no significant interaction of genotype with age and/or sex, suggesting that all mice decline similarly with age, regardless of AD carrier status, with female mice exhibiting the expected sexual dimorphism in muscle strength (Figure 5C).

**Figure 5:**
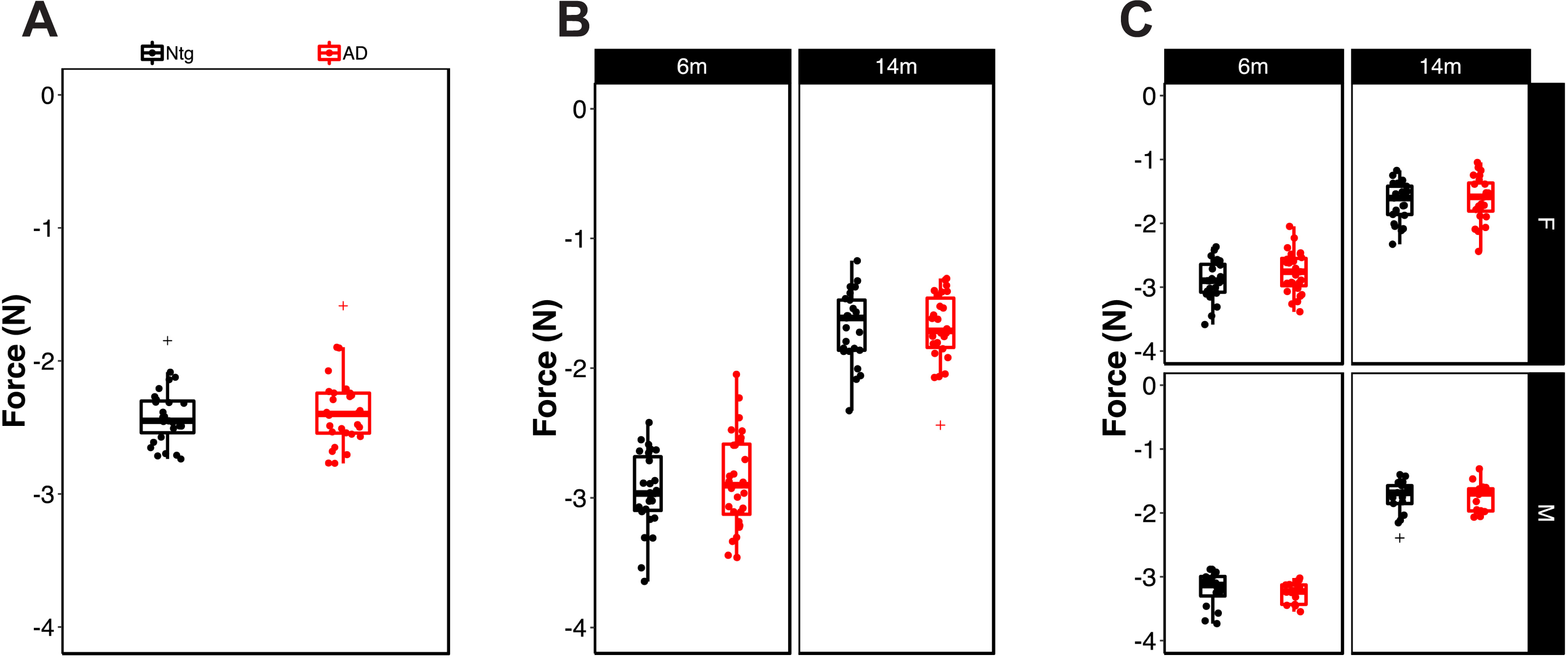
Age, but not sex or genotype, is associated with loss of grip strength. **A)** Grip strength in all four paws in Ntg- and AD-BXD strains. **B)** Grip strength stratified by age and **C)** by age and sex. Each point represents a strain average and statistical outliers are indicated by a + sign.

We determined the influence of genetic background on grip strength by assessing average grip strength for each strain at 6m and 14m (Figure 6A). As shown in Figure 6A and B, there is significant strain-dependent variation in grip strength in both Ntg and AD populations (Strain_Ntg_: F_(26, 363)_ = 3.02, p = 2.0 × 10^−6^; Strain_AD_: F(_27,474)_ = 4.2, p = 4.5 × 10^−11^) and a significant interaction between age and strain (Age*Strain_Ntg_: F_(24,363)_ = 2.6, p = 7.09 × 10^−5^; Age*Strain_AD_: F_(25,449)_ = 2.34, p = 0.0003). Consistent with our finding that there was no influence of genotype on grip strength, we found no significant interaction of genotype with strain (F_(26,839)_ = 0.37, p = 0.99). Taken together, these results suggest that age and genetic background profoundly impact muscle strength and decline and that this is not impacted by the presence of the AD transgene. As with the other sensorimotor traits assessed in this study, the correlation between grip strength in the Ntg-BXD and AD-BXD is not significant, although it is stronger than observed for narrow beam and inclined screen (*r* = 0.35, p = 0.09), suggesting that individual strains may perform more similarly at this task.

**Figure 6:**
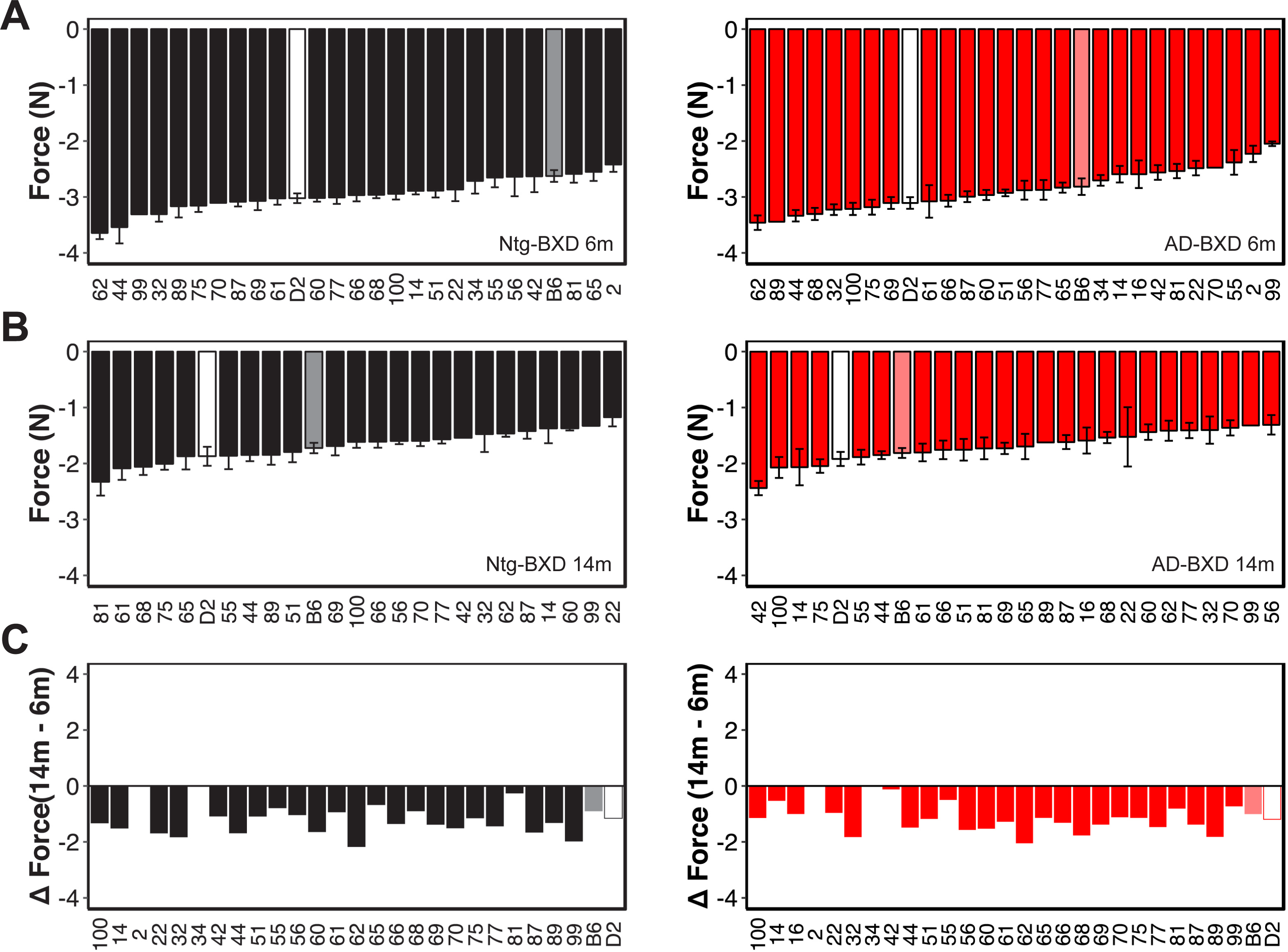
Genetic background influences grip strength in both Ntg- and AD-BXD strains. **A)** Mean grip strength in all four paws per strain in 6m old Ntg-BXD (*left*) and AD-BXD (*right*) strains. **B)** Mean grip strength in all four paws in 14m old Ntg-BXD (*left*) and AD-BXD (*right*) strains. **C)** Average-age related decline in four-paw grip strength in the Ntg-BXD (*left*) and AD-BXD (*right*) panels. Decline was calculated by subtracting grip strength at 6m of age from that measured at 14m of age. Data are presented as mean ± SEM. In all panels, number on the x-axis indicates the BXD strain used to generate each line.

## Discussion

### AD-BXD panel recapitulates the exacerbated age-related sensorimotor deficits associated with AD

Although dementia and cognitive decline are the hallmark diagnostic symptoms of Alzheimer’s dementia in humans, many AD patients also experience impairments in non-cognitive domains, including phenotypes associated with sensorimotor function such as coordination, balance, muscle strength, and proprioception (Albers et al., 2015; Buchman and Bennett, 2011; Buchman et al., 2009; Buchman et al., 2007; Portet et al., 2009; Tang et al., 2016; Voglein et al., 2019). In this study, we investigated the impact of naturally occurring genetic variation on age- and AD-related sensorimotor decline using our recently developed AD-BXD genetic reference panel (Neuner et al., 2019a). There was significant variation in balance, coordination, and vestibular function – both Ntg- and AD-BXD mice showed age-related declines in these phenotypes and the presence of familial AD mutations in the AD-BXD mice exacerbated this decline. There was a significant effect of sex on AD-related motor decline, with female AD-BXD mice exhibiting greater impairment on motor phenotypes than male AD-BXD mice. We found no relationship between motor function and decline in Ntg- and AD-BXD strains. Finally, although there was a pronounced effect of age on grip strength, we did not observe an effect of transgene on this phenotype.

### Influence of genetic background on sensorimotor traits and impact of the AD transgene on decline

We recently developed the first mouse model of genetic diversity in AD – the AD-BXDs and demonstrated that the inclusion of naturally occurring genetic variation introduces significant phenotypic variation in cognitive traits including long-term memory. Specifically, despite all strains carrying the same high-risk 5XFAD transgene, some strains exhibit resilience to cognitive impairment while others show increased susceptibility (Neuner et al., 2019a). We now extend that analysis to sensorimotor traits, including coordination, balance, vestibular function, and muscle strength. Similar to what we observed for cognitive abilities, there was a significant impact of genetic background on both baseline performance on motor tasks as well as both age- and AD-related decline. As with long-term memory, some strains exhibit resilience to age-related and/or AD-related impairments, exhibiting little decline with age, regardless of genotype. In contrast some strains even showed improvement with age, suggesting modifier genes exist that influence both the normal motor function-related aging process and AD-related decline in motor abilities.

Humans exhibit substantial variation in longevity and healthspan (Crimmins, 2015; Govindaraju et al., 2015; Sebastiani et al., 2017), including the development of frailty and physical impairment in old age (Buchman et al., 2009). Moreover, only a subset of AD patients exhibit motor-related impairments (Aggarwal et al., 2006; O’Keeffe et al., 1996; Portet et al., 2009; Voglein et al., 2019). The Ntg- and AD-BXD panel exhibits similar phenotypic variation, suggesting that these panels represent a key resource for investigating the influence of genetic variation on age- and AD-related frailty and motor decline. Heritability (*h*^*2*^) estimates for all three sensorimotor domains evaluated in this study range from 0.6 – 0.9 (Table 1) (Belknap, 1998), suggesting a strong influence of genetic background on these phenotypes. Thus, future studies will exploit the power of the AD-BXD panel as a mapping population to identify potential candidate modifier genes, which may represent novel targets for addressing sensorimotor decline in both normal aging and AD.

### AD-related motor decline is likely distinct from both AD-related cognitive decline and normal age-related decline

Frailty and motor decline are common in aging humans (Buchman et al., 2009), thus one possible explanation for the increased motor decline observed in both humans with AD and the AD-BXDs is that the AD disease process acts to accelerate ‘normal’ age-related motor decline (Albers et al., 2015; Buchman and Bennett, 2011). However, our results suggest that AD-related decline likely occurs via a distinct mechanism. In the two tasks that exhibited sensitivity to the AD transgene, narrow beam and incline screen, we found no correlation between the performance of the Ntg strains compared to the AD-BXD strains, suggesting that the underlying genetic mechanisms regulating motor impairment in these two models is distinct. There is similarly no correlation between sensorimotor and cognitive performance across the AD-BXD panel, including the Ntg-BXD strains (Neuner et al., 2019a), suggesting that AD-related motor impairment is distinct from both AD and age-related cognitive decline.

### Limitations of this study

Notably, although grip strength has been reported to be associated with AD risk in humans (Albers et al., 2015; Buchman and Bennett, 2011), we did not detect an effect of AD genotype on grip strength in the AD-BXD panel at either age tested, although there was a pronounced effect of age on muscle strength in both Ntg- and AD-BXD mice. However, there are several reports in human cohorts that indicate that lower extremity dysfunction is more closely associated with mild cognitive impairment (MCI) and AD than upper limb function (Aggarwal et al., 2006; Boyle et al., 2007; Eggermont et al., 2010; Goldman et al., 1999; Kluger et al., 1997). Our assessment of grip strength measured force exerted by all four limbs in each mouse and thus cannot distinguish between differential impairments between the forelimbs and hindlimbs in mice, although the narrow beam and inclined screen tasks would likely be sensitive to limb-specific decline. Additionally, it is important to note that expression of the 5XFAD transgene used to generate the AD-BXD reference panel is driven by a *Thy-1* promoter and likely does not fully recapitulate the endogenous pattern or amyloid pathology or presenilin expression in the periphery. This limitation can be overcome by the use of a ‘knockin’ model in which human mutations in *App* and *Psen1* are controlled by the endogenous promoters for both genes to produce a more translationally-relevant organism-wide expression pattern in both the CNS and peripheral tissues.

In human AD patients, motor impairment has been reported to occur prior to the onset of cognitive symptoms (Aggarwal et al., 2006; Albers et al., 2015; Buchman and Bennett, 2011; Voglein et al., 2019), but in the AD-BXD panel we did not observe significant impairment on any of the sensorimotor tasks used in this study at the earliest age measured (6m). While we cannot exclude the possibility that motor decline does not precede cognitive deficits in the AD-BXDs, a limitation of our study design is that motor function was measured at only two ages, 6m and 14m. The average age at onset for cognitive impairment in the AD-BXD panel is ∼10m (Neuner et al., 2019a), thus motor performance at 6m of age may simply be too early of an age to detect a preclinical decline in sensorimotor function in the AD-BXDs. In support of this hypothesis, although not significant (p= 0.08), we do see a trend for worse narrow beam performance in the AD-BXDs compared to Ntg-BXDs at 6m of age (Figure 1B).

Some caution is warranted in the interpretation of narrow beam data when considering the behavior of mice during this task. We report strains with averages of latency > 120s (Figure 1C), suggesting that some strains have a large number of animals that simply did not move while on the beam or failed to remain on the beam. As a result, in some cases, an increase in time to cross the beam may not necessarily be due to an impairment of sensorimotor performance, but rather another behavioral output such as anxiety that influences performance on these tasks.

## Conclusions

The results presented here provide further evidence for the translational utility of our novel AD-BXD panel, which we now demonstrate also exhibits sensorimotor decline similar to that reported in human AD patients. Moreover, this decline is strongly modified by genetic background and exhibits a high degree of heritability, consistent with the human disease. Thus the AD-BXD panel represents a novel tool to investigate sensorimotor deficits in normal aging and AD and facilitate the discovery of novel targets to address these deficits. Our finding that there is no correlation between motor and cognitive phenotypes in either the AD strains or the normal aging controls indicates that motor decline likely occurs via a mechanism distinct from cognitive decline in both AD and normal aging individuals. Future work will incorporate additional strains for genome-wide mapping to identify potential modifier genes and omics analysis of CNS regions associated with sensorimotor tasks.

## Methods

### Mice

Generation of the AD-BXD panel was described in detail in Neuner et al 2019. Briefly, female congenic C57Bl/6J mice hemizygous for the 5XFAD transgene (Oakley et al., 2006) (MMRRC Stock No: 34848-JAX) were crossed to male mice from strains selected from the BXD genetic reference panel (Peirce et al., 2004). The resulting F1 offspring are isogenic recombinant inbred mice carrying one maternally derived *B* allele and one paternally derived *B* or *D* allele at each genomic locus. Furthermore, ∼50% of F1 mice carry the 5XFAD transgene (termed AD-BXD), while the other ∼50% are nontransgenic littermate controls (termed Ntg-BXDs). Mice from a total of 27 Ntg- and AD-BXD (27 female, 18 male) strains were used in this study; mice were genotyped for the 5XFAD transgene by either the Transgenic Genotyping Service at The Jackson Laboratory or Transnetyx, Inc. All mice were fed a standard laboratory mouse chow (Teklad 8604) and both food and water were available *ad libitum*. All mice were kept on a 12:12 light cycle and were phenotyped as described in Neuner, et al 2019 (Neuner et al., 2019a).

All experiments involving mice were performed at the University of Tennessee Health Science Center and were approved by the Institutional Care and Use Committee at that institution and carried out in accordance with the standards of the Association for the Assessment and Accreditation of Laboratory Animal Care (AAALAC) and the NIH Guide for the Care and Use of Laboratory Animals.

### Sensorimotor phenotyping

Sensorimotor function was assessed at 6 and 14 months of age in the Ntg-BXD and AD-BXD panels. A total of 894 mice (575 female, 319 male) 27 AD- and Ntg-BXD strains were included in this study. The sensorimotor tasks used were narrow beam, negative geotaxis (incline screen), and grip strength (Neuner et al., 2019a).

#### Narrow beam

Briefly, the narrow beam task was used to assess motor and balance coordination by placing each mouse in the center of a 1 meter long beam 12 mm wide that was elevated 50 cm above a table surface (Maze Engineers, Boston MA). The time taken in seconds for each mouse to cross the beam to a safe platform on either side was recorded. A maximum time limit of 180 s was imposed; if a mouse fell from the beam, the maximum time of 180 s was given. The average of three trials for the task was used to assess the performance for each mouse and the average used for statistical analysis.

#### Incline screen

The incline screen task was used to assess vestibular and/or proprioceptive function. Each mouse was placed nose-down in the center of a wire mesh grid (1 cm × 1 cm) positioned at a 45° angle (Harvard Apparatus). The time taken for a mouse to reorient itself with its nose facing upwards (negative geotaxis) was recorded; the average of three trials for each mouse was used for analysis. As with narrow beam, a 180 second maximum time limit for righting was used.

#### Grip strength

Muscle strength was measured using a standard grip strength meter. Each mouse was placed vertically on the wire grid of the apparatus (Columbus Instruments) with all four paws grasping the grid and then gently pulled away from the grid by the base of its tail until its grip was released to measure the force exerted. The average of three trials for each mouse was recorded and used for analysis.

All data and statistical analysis was performed using R. A univariate ANOVA was used for each sensorimotor task using genotype, age, sex, and strain as fixed factors. Data are reported as mean ± standard error of the mean in both the main text and figure legends. A Pearson’s correlation test was used to assess the relationship of decline between the Ntg-BXD and AD-BXD on these tasks. CFA and CFM measures used for correlations in Supplemental figures 2 and 3 are from (Neuner et al., 2019a). All raw data used in this study are available through the Synapse AMP-AD Knowledge Portal (https://www.synapse.org/#!Synapse:syn17016211). Strain averaged behavioral data has also been deposited at GeneNetwork.org. Heritability estimates for the sensorimotor phenotypic traits in the AD- and Ntg-BXDs were determined by calculating the ratio of between-strain variance to total sample variance (variance due to both genetic and technical/environmental factors) according to (Belknap, 1998).

## ACKNOWLEDGMENTS

This study is part of the National Institute on Aging Resilience-AD program and is supported through the NIA grant award R01AG057914 to C.C.K. This work was also supported by the BrightFocus Foundation (A2016397S to C.C.K.), the National Institute on Aging (R01AG054180 to C.C.K.; F31AG050357 to S.M.N.; RF1AG059778 to K.M.S.O. and C.C.K.), and the National Institute of Diabetes and Digestive and Kidney Diseases (R01DK102918 to K.M.S.O.). Additional support was provided by the Evnin family and the University of Tennessee Health Science Center Neuroscience Institute. The authors thank Dr. Lynda Wilmott and Thomas Shapaker for collection of behavioral data. The authors would also like to thank Dr. Rob Williams for thoughtful input on the project design.

## AUTHOR CONTRIBUTIONS

K.M.S.O., S.M.N. and C.C.K. conceived of the experiments. S.M.N. conducted the behavioral experiments. K.M.S.O., A.R.O and C.C.K. designed the analyses. A.R.O. and K.M.S.O. performed the data analysis. A.R.O., C.C.K., S.M.N., A.R.D, and K.M.S.O. contributed to the interpretation of results. K.M.S.O. wrote the manuscript. All authors approved of the final manuscript.

---

**Supplemental figure 1:**
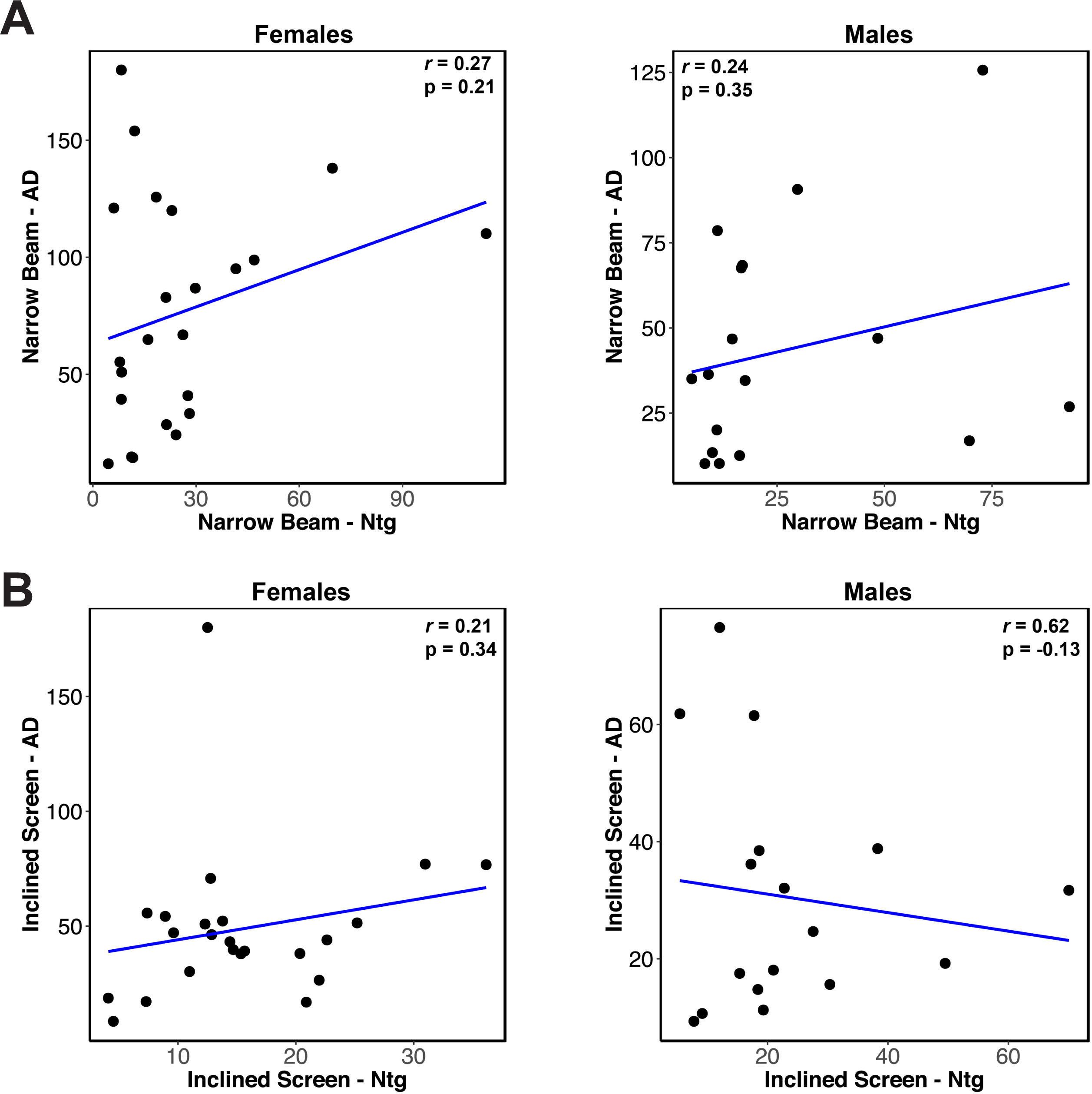
Lack of correlation for motor-related phenotypes between genotypes suggests AD-related motor decline is distinct from decline related to normal aging. **A)** Scatterplot of narrow beam performance at 14m of age in female (*left*) and male (*right*) Ntg- and AD-BXD strains. **B)** Scatterplot of inclined screen performance at 14m of age in female (*left*) and male (*right*) Ntg- and AD-BXD strains.

**Supplemental figure 2:**
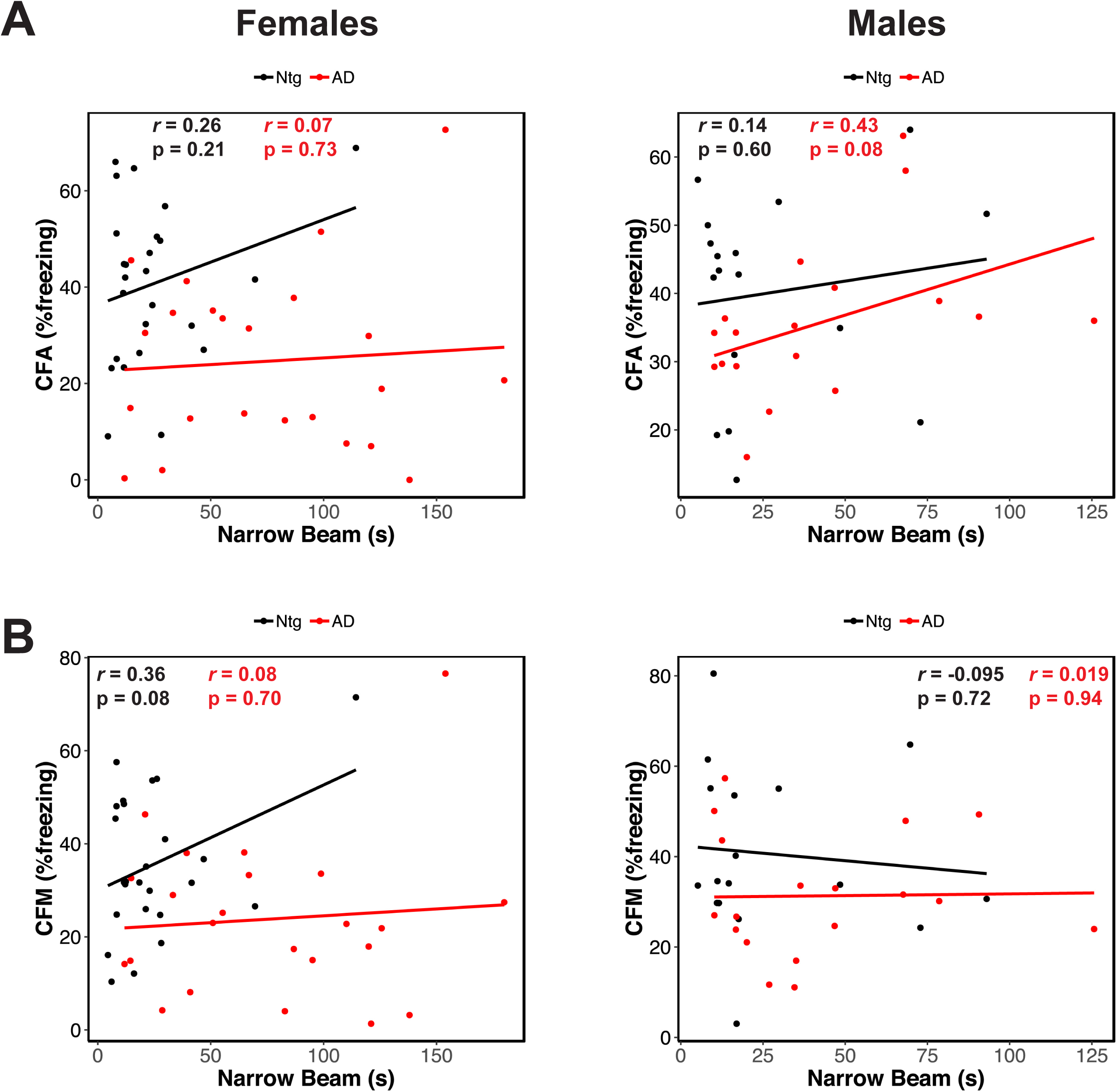
Lack of correlation between narrow beam performance and cognitive phenotypes. Narrow beam performance in 14m old female (*left*) and male (*right*) plotted against contextual fear acquisition (CFA) **(A)** and contextual fear memory (CFM) **(B)**. Each point represents a strain mean. Black = Ntg-BXD; red = AD-BXD.

**Supplemental figure 3:**
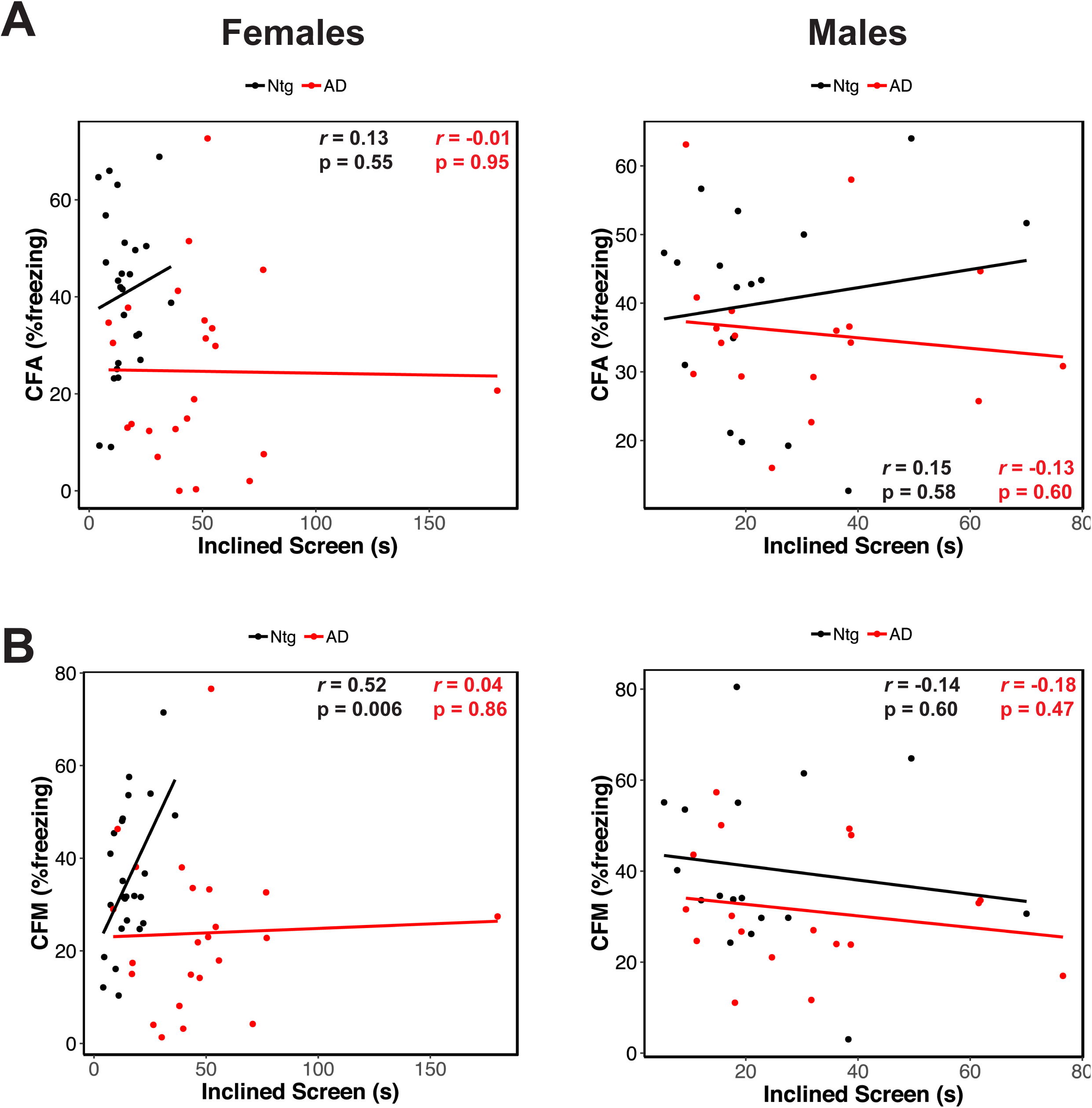
Lack of correlation between inclined screen performance and cognitive phenotypes. Inclined screen performance in 14m old female (*left*) and male (*right*) plotted against contextual fear acquisition (CFA) **(A)** and contextual fear memory (CFM) **(B)**. Each point represents a strain mean. Black = Ntg-BXD; red = AD-BXD.

